# Length control emerges from cytoskeletal network geometry

**DOI:** 10.1101/2023.11.28.569063

**Authors:** Shane G. McInally, Alexander J.B. Reading, Aldric Rosario, Predrag R. Jelenkovic, Bruce L. Goode, Jane Kondev

**Affiliations:** Department of Biology and Biotechnology, Worcester Polytechnic Institute, Worcester, MA, 01609, USA; Department of Physics, Brandeis University, Waltham, MA, 02454, USA; Department of Biology, Brandeis University, Waltham, MA, 02454, USA; Department of Electrical Engineering, Columbia University, New York, NY 10027

**Keywords:** Cytoskeleton, Size control, Biological scaling, Emergence

## Abstract

Many cytoskeletal networks consist of individual filaments that are organized into elaborate higher order structures. While it is appreciated that the size and architecture of these networks are critical for their biological functions, much of the work investigating control over their assembly has focused on mechanisms that regulate the turnover of individual filaments through size-dependent feedback. Here, we propose a very different, feedback-independent mechanism to explain how yeast cells control the length of their actin cables. Our findings, supported by quantitative cell imaging and mathematical modeling, indicate that actin cable length control is an emergent property that arises from the cross-linked and bundled organization of the filaments within the cable. Using this model, we further dissect the mechanisms that allow cables to grow longer in larger cells, and propose that cell length-dependent tuning of formin activity allows cells to scale cable length with cell length. This mechanism is a significant departure from prior models of cytoskeletal filament length control and presents a new paradigm to consider how cells control the size, shape, and dynamics of higher order cytoskeletal structures.

**Significance Statement:** Cells control the sizes of their cytoskeletal networks to ensure that these structures can efficiently perform their cellular functions. Until now, this ability has been attributed to molecular feedback mechanisms that control the rates at which individual filaments are assembled and disassembled. We find that size control of cytoskeletal networks does not require this type of feedback and can instead be encoded through the physical arrangement of the filaments within that network. These findings have important implications for understanding how the underlying geometry of higher order cytoskeletal networks contributes to cellular control over these structures.

## Introduction

Cells possess the remarkable ability to control the size, shape, and dynamics of their intracellular parts (1–3). This behavior is important for promoting proper organelle function and has been observed for many membrane-bound and cytoskeletal organelles found in diverse eukaryotic cells. Further, it suggests that the ability of a cell to govern the geometric properties of its intracellular structures is a fundamental property of living systems.

Cytoskeletal filaments are popular and convenient models used to study the mechanisms that control the size of intracellular structures because their size can be represented by a single dimension, their length. Regardless of their molecular composition (e.g., actin or tubulin), these polymers grow by the addition of molecular building blocks and shrink by their removal. Thus, experimental and theoretical studies of length control aim to identify the nature of the feedback that controls the rates of subunit addition and removal, which allows these filaments to be assembled and maintained at a steady-state length (4). Different mechanisms have been proposed to explain how cytoskeletal structures (e.g., mitotic spindles, cilia, and actin cables) are assembled and maintained at defined lengths, including: limiting-pool models, balance-point models, molecular rulers, antenna models, and concentration gradients (5–10). While each of these mechanisms involves distinct molecular details, they all require a control mechanism that tunes either the assembly rate, the disassembly rate, or both rates in a length-dependent manner. While this level of abstraction is suitable for individual cytoskeletal filaments, it is unclear how well these types of models explain size control of the many higher order cytoskeletal structures found in cells, which have elaborate filamentous architectures, such as cilia/flagella, stereocilia, lamellipodia, and filopodia.

Cytoskeletal networks found in nature are typically composed of many individual filaments that are organized into higher order structures. The specific architectures of these larger, composite structures are crucial for their biological functions (e.g., in phagocytosis, cell motility, and pathogenesis), yet much of the work investigating how these structures are assembled and regulated has focused on the mechanisms that control the turnover of individual filaments. To gain a better understanding of how these higher order structures are controlled by the cell, we need to consider the architecture and geometry of these structures and how the arrangement of filaments within these structures contributes to emergent properties of these higher order networks.

Here, we address this question using yeast actin cables as a model. Each actin cable in a yeast cell is a bundle comprised of many short, overlapping actin filaments polymerized by formins (11). In the budding yeast *S. cerevisiae*, cables are assembled by two genetically redundant formins, which localize during bud growth to the bud tip (Bni1) and bud neck (Bnr1) (12–14). The cables polymerized by Bni1 and Bnr1 are polarized structures, with their barbed ends oriented toward the bud tip and neck, respectively. This property enables them to serve as railways for essential myosin-based transport of secretory vesicles and organelles to the growing bud cell (11, 15). It is thought that controlling actin cable length promotes efficient intracellular transport and therefore polarized growth in these cells (9, 16–18). In support of this hypothesis, we have recently shown that yeast actin cables grow so that their length closely matches the length of the mother cell in which they are assembled (19). We found that the scaling of cable length with cell length is conferred through control over their assembly - initially cables grow fast, but as they grow longer and approach the back of the cell their rate of growth steadily slows down, or decelerates. Ultimately, cable growth stops when the length of the cable matches the length of the cell. In addition, we showed that this cable deceleration behavior was different in smaller versus larger cells. This suggests that cable growth is tuned in a cell length-dependent manner, but the underlying mechanism has remained unclear.

Here, we present a new mathematical model of cable length control that explores how the specific geometry and architecture of a cable can enable length control. This model for cytoskeletal length control is a significant departure from previous length control models because there is no length-dependent molecular feedback mechanism that tunes the rates of assembly or disassembly. Instead, the control over cable length naturally emerges from the geometric arrangement of the filaments within the network.

## Results

### Actin cables undergo length-dependent tapering

To date, actin cables have been thought of as one-dimensional, linear structures (9, 18, 19). Therefore, prior length control studies have treated the cable as having a single barbed end at which actin monomers are added, and a single pointed end at which actin monomers are removed. However, it has been shown that cables are composed of many shorter, overlapping actin filaments bundled together by actin crosslinkers (20). Therefore, we were interested in determining whether the architecture of the cable could provide insights into how its length is controlled (21– 23).

We started by asking whether the width of cables is uniform along their lengths. To address this, we fixed and stained wildtype haploid budding yeast cells with fluorescently labelled phalloidin and imaged them using super-resolution microscopy. From these images, we traced the entire length of the cables that could be clearly tracked in mother cells (i.e., those that do not intersect with other cables or actin patches) from their origin at the bud neck to their terminal end in the mother cell (Figure 1A). We measured the fluorescence intensity along the entire length of the cable, and took this to be proportional to cable density or thickness. We found that the density of F-actin in cables was not uniform, but instead tapers as cables get longer (Figure 1B). Specifically, F-actin density was greatest in the region closest to the bud neck, where formin-mediated cable assembly takes place, and progressively decreased with cable length. Further, the cable thickness profile was well fit by a single exponential with a decay length of 1.54±0.08 µm (all reported values represent mean ± 95% CI, unless otherwise indicated).

**Figure 1:**
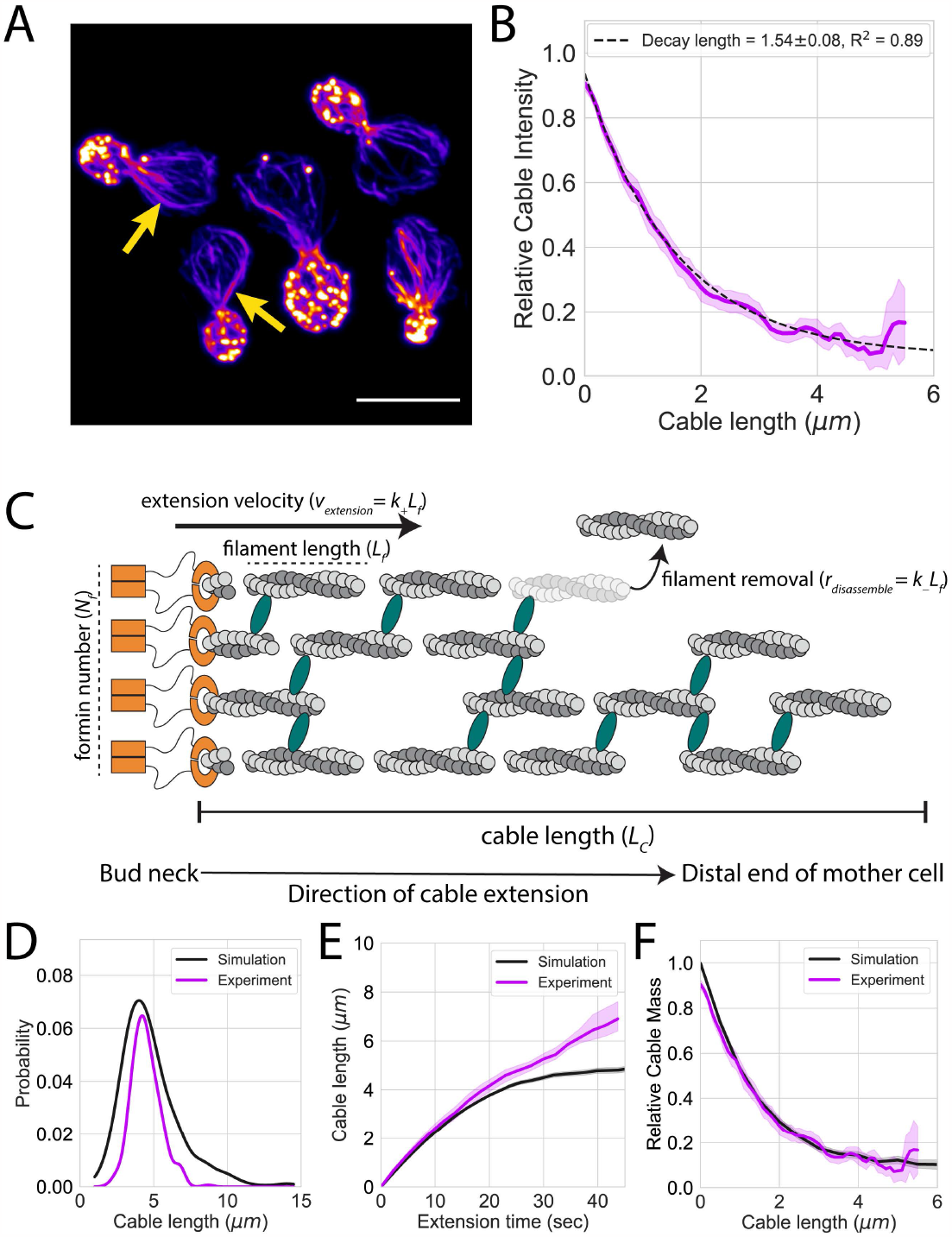
Two-dimensional model of cable length control. (A) Representative maximum intensity projection images of haploid yeast cells fixed and stained with labeled-phalloidin. Arrows indicate single actin cables that clearly display their tapered shape. Scale bar, 5µm. (B) Relative actin cable fluorescence intensity measured in three independent experiments. Solid magenta line and shading, mean and 95% confidence interval for all three experiments (n=47 cables). Tapering profile decay length (±95% CI) was determined by fitting the profile to a single exponential. (C) Schematic of the two-dimensional model of actin cable length control. Multiple formins (orange, *N*_*f*_) simultaneously assemble short actin filaments with a characteristic length (*L*_*f*_) at a constant rate (*k*_+_). These filaments are crosslinked and bundled (green ellipses) with neighboring filaments to form the cable and continue to extend into the cell at the same rate at which filaments are assembled by formins (*v*_*extension*_ = *k*_+_*L*_*f*_). Each filament has an independent probability of being targeted for removal (*r*_*disassemble*_ = *k*_−_*L*_*f*_) from the cable. Thus, the length of the cable (*L*_*c*_) is the distance from the site of assembly to the distal tip of the longest surviving filament in the cable. (D-F) Results obtained from simulations (solid black lines) compared with experimental measurements of cable length (D), cable extension rate (E), and cable tapering (F). The parameters used for these 1,000 independent simulations were, *k*_+_ = 0.50 sec^-1^, *k*_−_= 0.16 sec^-1^, *L*_*f*_ = 500nm, *N*_*f*_ = 4 formins. Solid lines and shading indicate mean and 95% confidence interval, respectively.

### Two-dimensional model of cable length control

The tapering of F-actin density we observed in actin cables was reminiscent of tapering previously reported for other types of actin networks (e.g., *Listeria* comet tails and fish keratocyte lamellipodial fragments) (24–26). This prompted us to consider whether related mechanisms may explain how the structure and length of actin cables are regulated. To test this idea, we developed a mathematical model of cable length control (Figure 1C), in which multiple formin molecules (*N*_*f*_) are localized at the bud neck and produce actin filaments of a fixed length (*L*_*f*_) at a constant rate (*k*_+_). As these filaments are assembled, they are incorporated into the cable bundle by crosslinkers. As a result of polymerization and crosslinking, the entire bundle collectively grows as a single unit, extending into the mother cell at a constant velocity (*v*_*extension*_ = *k*_+_*L*_*f*_), which is equivalent to the number of actin monomers that are added to the growing cable the formins at the bud neck. Once incorporated into the growing bundle, each filament has an independent probability of being targeted for removal through an unspecified disassembly mechanism. Because each of these filaments has a fixed length (*L*_*f*_), the rate at which monomers are removed from the cable is constant (*r*_*disassemble*_ = *k*_−_*L*_*f*_). Thus, the entire length of a cable (*L*_*c*_) is equal to the distance between its site of assembly (the bud neck) and its distal end, defined by the last surviving filament within the bundle. Importantly, none of these parameters have an inherent length dependence, and therefore all parameters in this model are constants.

To derive estimates for the parameters in our model, we referred to our prior study of cable length control (19), in which we determined that the average length of cables in haploid budding yeast was 4.48 ± 0 .98 μm. We also used linear regression to measure the extension velocity of cables (i.e., the slope of the initial linear phase of cable growth) from our prior measurements of cable extension rates in haploid cells (*v*_*extension*_ = 0 .25 ± 0 .0 2 μm/sec, Supplemental Figure 1A). To estimate the remaining unmeasured parameters in our model, we used the following mathematical relationship that describes the mean length of a bundle of filaments:

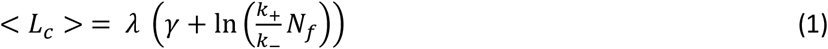

where < *L*_*c*_ > is the mean cable length, γ ≈ 0.577 (i.e., the Euler-Mascheroni constant), and *N*_*f*_ is the number of formins assembling a single actin cable; for derivation see Supplemental Text. Importantly, λ is the cable tapering profile in Figure 1B, and can be related to the model parameters through the thickness decay constant, defined as:

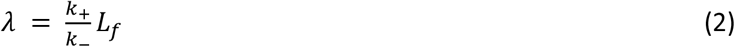

Using Equation 2 with our measurements of the extension velocity (*v*_*extension*_) and the tapering decay profile (λ) we estimate *k*_−_ = 0.16±0.01 sec^−1^ (mean ± SD).

While we were unable to compute *N*_*f*_ and *L*_*f*_ without direct measurements of at least one of these parameters, a prior electron microscopy study of actin cables in *S. pombe* found that the average length of these filaments was 0.49 ± 0.26 μm (mean ± SD) (20). We used these measurements to estimate *L*_*f*_∼0.5μm and, with Equation 1, estimate that four formins (*N*_*f*_ ∼4 formins) cooperate to assemble a single cable. Importantly, rewriting Equation 1 using Equation 2 as:

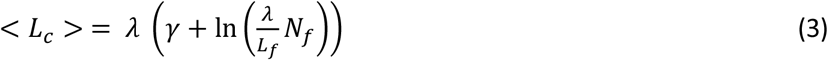

shows that mean cable length depends on the ratio of the number of formins to filament length (i.e. *N*_*f*_ /*L*_*f*_), indicating that other values for these parameters can generate cables with the same average length when λ is held constant (Supplemental Figure 2A).

Next, we conducted computational simulations using the parameters estimated above (*k*_+_ = 0.50 sec^−1^, *k*_−_ = 0.16sec^−1^, *L*_*f*_ = 500 nm, *N*_*f*_ = 4 formins) and found that this model can assemble actin cables that resemble those observed in vivo. Remarkably, our model produced cables that exhibit a peaked distribution of lengths, decelerated growth, and tapered actin density profiles (Figure 1D-F, black lines), despite the absence of any length-dependent parameters. Next, we directly compared the results of these simulations with our experimental measurements (Figure 1D-F, magenta lines) and found that this model can adequately recapitulate our experimental data without the use of any fitted parameters. We further validated our simulations by comparing these results with the analytical solutions for these cable behaviors (Supplemental Figure 2B-D).

### Cable extension velocity is independent of cell size

Next, we wanted to determine which parameters in our model may be tuned in a cell length-dependent manner to permit the previously observed scaling of cable length with cell length (19). First, we considered whether the extension velocity may be cell length dependent. To determine how extension velocity changes as a function of cell size we referred to our prior quantification of cable extension rates from temperature-sensitive *cdc28-13*^*ts*^ cells. At the permissive temperature, *cdc28-13*^*ts*^ cells are similar in size to wildtype haploid budding yeast; however, their size increases when grown at the non-permissive temperature (19, 27, 28). In our prior study, we quantified cable extension rates from these enlarged cells by tracking the tips of cables marked with the fluorescent cable reporter Abp140-GFP^Envy^. Here, we reanalyzed these measurements by using linear regression to compare the extension velocity (i.e., the slope of the initial linear phase of actin cable growth) in induced and uninduced *cdc28-13*^*ts*^ cells. We found that despite the nearly 2-fold difference in cell length, the initial extension velocity was not significantly different (*v*_*extension, uninduced*_ = 0 .22 ± 0 .0 2 μm/sec, *v*_*extension, uninduced*_ = 0 .24 ± 0 .0 2 μm/sec; p = 0.23)(Supplemental Figure 1B). Thus, cable initial extension velocity is independent of cell size and does not likely contribute to the scaling of cable length with cell length.

### The amount of formin at the bud neck scales with cell length

Next, we considered whether differences in the density or organization of formin molecules at the bud neck (Bnr1) might contribute to the scaling of cable length with cell length. To determine whether the amount of Bnr1 at the bud neck changes in cells of different size, we differentially tagged Bnr1 with GFP^Envy^ and Cdc3 (a component of the septin collar at the bud neck) with mCherry in *cdc28-13*^*ts*^ cells (Figure 2A). We grew the cells for either 0, 4, or 8 hours at the non-permissive temperature to induce different changes in cell size, and then returned cells to the permissive temperature to allow polarized growth for one hour. Next, we mixed approximately equal numbers of cells of the three different sizes and performed live imaging on the cell populations using spinning disk confocal microscopy. We used the Cdc3-mCherry channel to generate segmentation masks of the bud neck, and within this mask measured the total fluorescence intensity of Bnr1-GFP^Envy^ at the bud neck. From the same images, we also measured the distance from the bud neck to the rear of the mother cell.

**Figure 2:**
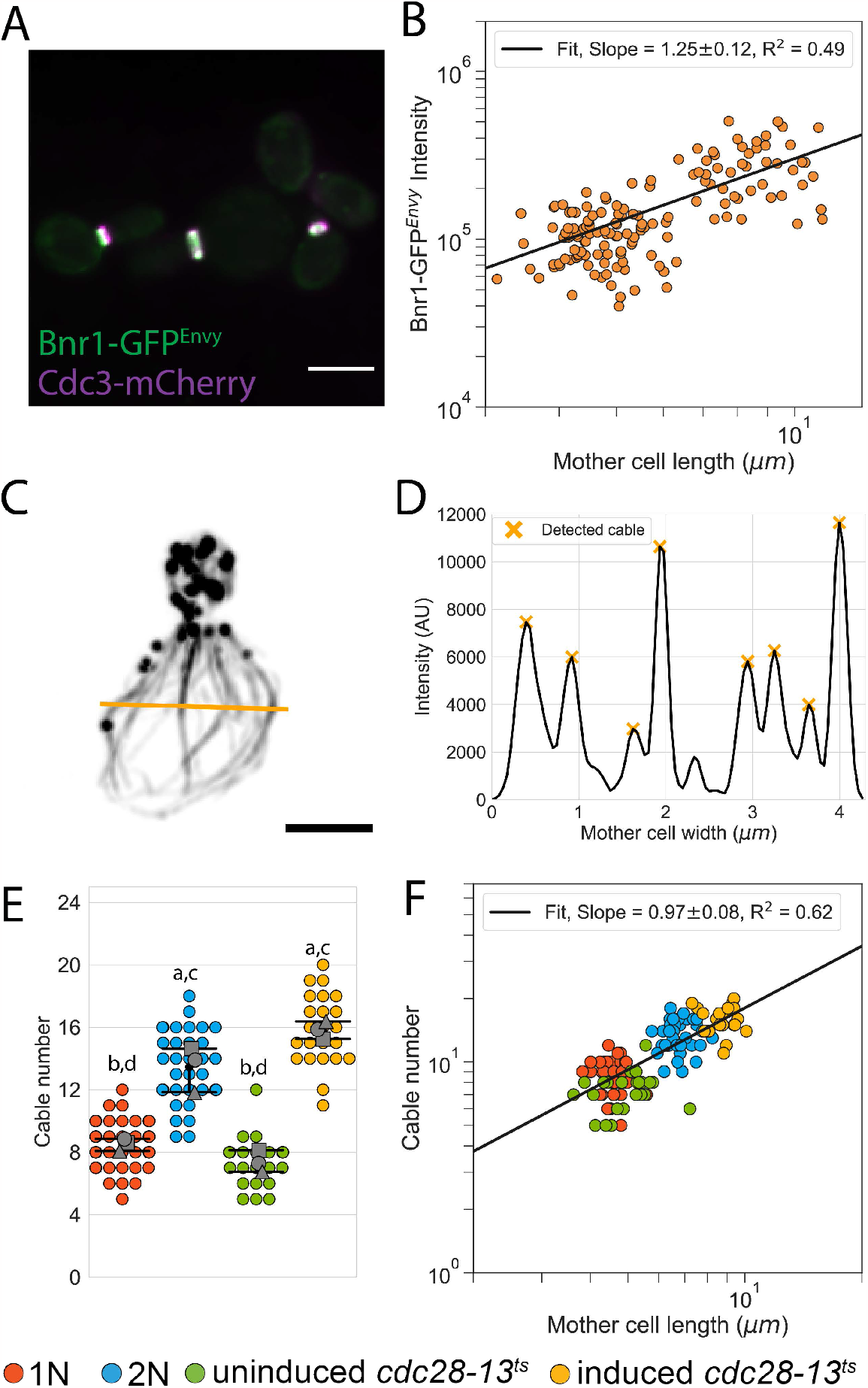
The amount of Bnr1 formin at the bud neck and the number of actin cables in a cell scale with cell length. (A) Representative maximum intensity projection image of *cdc28-13*^*ts*^ cells grown to different sizes while expressing fluorescently labeled Bnr1 (Bnr1-GFP^Envy^) and Cdc3 (Cdc3-mCherry). Scale bar, 5µm. (B) Amount of Bnr1-GFP^Envy^ localized to the bud neck of *cdc28-13*^*ts*^ cells grown to different sizes plotted against mother cell length on a double logarithmic plot and fit using the power-law. Bnr1-GFP^Envy^ was measured in three independent experiments (n=148 cells). (C) Representative maximum intensity projection images of a haploid yeast cell fixed and stained with labeled-phalloidin. Scale bar, 2µm. Yellow bar indicates the ROI position used to generate the line scan profile (D) used for automated peak detection (orange X’s indicate detected actin cables). (E) The number of actin cables measured from haploid (red), diploid (blue), uninduced *cdc28-13*^*ts*^ (green), and induced *cdc28-13*^*ts*^ (yellow) cells fixed and stained with labeled-phalloidin. Each data point represents an individual cell. Larger symbols represent the mean from each of the three independent experiments (n=119 cells). Error bars indicate 95% confidence intervals. Statistical significance determined by students t-test. Significant differences (p≤0.05) indicated for comparisons with haploid (‘a’), diploid (‘b’), uninduced cdc28-13ts (‘c’), and induced cdc28-13ts (‘d’). (F) Actin cable number plotted against mother cell length on a double logarithmic plot and fit using the power-law.

To determine whether the amount of Bnr1-GFP^Envy^ at the bud neck changes as a function of cell length, we analyzed the data on a double logarithmic plot, which revealed a linear scaling relation between the amount of Bnr1 at the bud neck and cell length (Figure 2B). To determine the nature of this scaling relation, we fit the data using the power law (*y* = *Ax*^*a*^), where *a* is the scaling exponent that describes the relationship between the two measured quantities, cell length and cable number (3). We found that the scaling exponent was slightly hyperallometric (*a*_*formin*_ = 1.25 ± 0 .11, R^2^ = 0.49), indicating that a greater amount of formin localized to the bud neck in larger cells compared to smaller cells.

### The number of actin cables in the mother cell scales with cell length

Our observations above prompted us to next ask whether larger cells, which have higher levels of Bnr1 at the bud neck, might assemble thicker cables and/or an increased number of cables. To quantify the number of cables in the mother cell compartment of cells of different size, we used line scans drawn across the equator of haploid, diploid, and *cdc28-13*^*ts*^ temperature-sensitive cells fixed and stained with fluorescently labelled phalloidin (Figure 2C). Diploid yeast cells have ∼2-fold increase in volume compared to haploid cells, and *cdc28-13*^*ts*^ cells grown at the non-permissive temperature for eight hours have a ∼5-fold increase in cell volume (19, 29). Next, we used automated fluorescent peak detection from the line scans to quantify the number of cables in the mother cell compartment (Figure 2D). We also measured the length of the mother cell (i.e., the distance from the bud neck to the rear of the mother cell) in each cell.

We found that the mean number of cables was 9 ± 2 in haploid cells and 13 ± 3 in diploid cells. Additionally, the mean number of cables in *cdc28-13*^*ts*^ cells grown at the permissive temperature was 7 ± 2, while the mean number of cables in *cdc28-13*^*ts*^ cells grown at the restrictive temperature was 16 ± 3 (Figure 2E). We performed a power law analysis to compare how the number of cables changes as a function of cell size, and found that there is an isometric scaling relation (*a*_*cable number*_ = 0 .97 ± 0 .0 7, R^2^ = 0 .62) between the number of cables and the length of the cell (Figure 2F).

### Actin cables taper in a cell length-dependent manner

Thus far, our data suggest that larger cells have higher levels of formin molecules at the site of cable assembly; however, instead utilizing these increased levels of formins to assemble thicker cables, they assemble more cables. To explicitly test whether cables are thicker in larger cells compared to smaller cells, we compared cable tapering profiles from uninduced and induced *cdc28-13*^*ts*^ cells, which were fixed and stained with fluorescently labelled phalloidin. To control for possible differences in staining efficiency, we mixed approximately equivalent amounts of uninduced and induced *cdc28-13*^*ts*^ cells and then simultaneously fixed, stained, and imaged them using super-resolution microscopy (Figure 3A). For each cell in the population, we measured the fluorescence intensity along the length of its cables and the length of the mother cell. To distinguish between the uninduced and induced *cdc28-13*^*ts*^ cells, we used mother cell length to sort cells into bins containing either ‘small’ or ‘large’ cells. To validate this binning strategy, we plotted the cable lengths we measured from these cells and found that the mean cable length in each bin was consistent with our previous measurements (*L*_*cable, small*_ = 4.1 ± 0 .3μm, *L*_*cable, large*_ = 7.3 ± 0 .8μm) (Figure 3B)(19).

**Figure 3:**
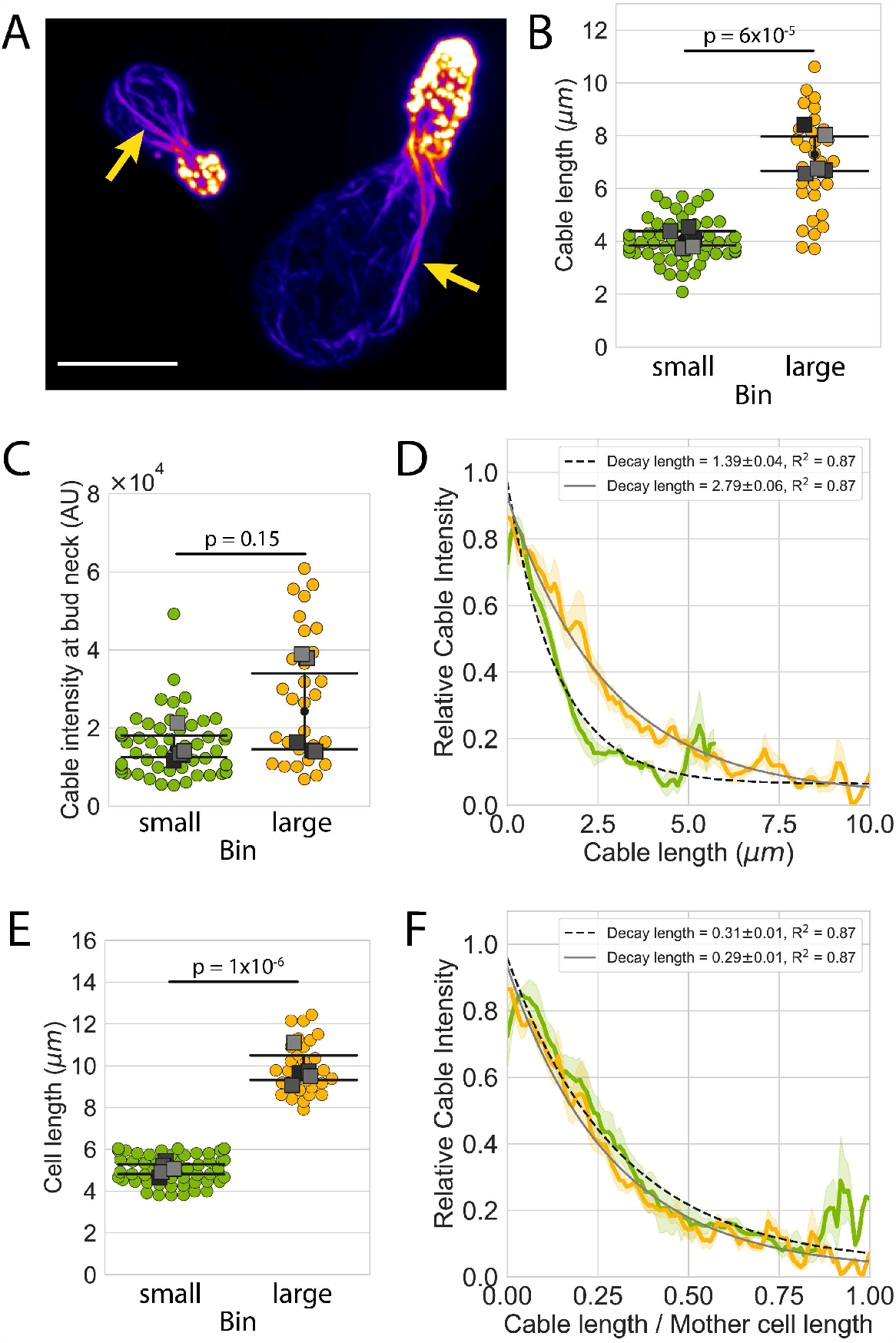
Actin cable tapering is cell length dependent. (A) Representative maximum intensity projection images of small (left) and large (right) *cdc28-13*^*ts*^ cells fixed and stained with labeled-phalloidin. Arrows indicate single actin cables that clearly display their tapered shape. Scale bar, 5µm. (B) Actin cable length and (C) actin cable fluorescence intensity in the bud neck region measured from mixed populations of uninduced and induced *cdc28-13*^*ts*^ cells. Cells were binned based on cell length (E), small cells are indicated in green while large cells are indicated in yellow. Each data point represents an individual cable. Larger symbols represent the mean from each experiment. Error bars indicate 95% confidence intervals. Statistical significance determined by students t-test. (D) Relative actin cable fluorescence intensity plotted against cable length, and (F) relative actin cable fluorescence intensity plotted against the ratio of cable length/cell length. Solid lines and shading, mean and 95% confidence interval. Tapering profile decay lengths (±95% CI) were determined by fitting each profile to a single exponential. All data were generated from five independent experiments (n=84 cables).

We first compared the cable fluorescence intensity at the region closest to the bud neck (i.e., the region where new filaments are added to the cable) and found that there was no statistically significant difference between these bins (Figure 3C). These findings indicate that the number of formins incorporating new actin filaments into a single cable is likely similar in cells of different size, and therefore does not contribute to the scaling of cable length with cell length.

We next wanted to determine whether differences in how filaments are removed from the cable bundle may contribute to the scaling of cable length with cell length. To test this, we measured the decay length (λ) from the actin tapering profiles for each bin, as this measurement directly reflects the rates at which filaments are added and removed from the bundle (see Equation 2). Comparing the decay length (λ) from the actin tapering profiles for each bin revealed that the decay length was ∼2-fold greater in larger compared to smaller cells (λ_*small*_ = 1.39 ± 0 .0 4 μm, λ_*large*_ = 2.79 ± 0 .0 6 μm) (Figure 3D). We also noted that the ratio of decay lengths between bins was similar to the ratio of average cell length between bins (*L*_*cell, large*_/*L*_*cell, small*_ = 2.0 ± 0 .3, λ_*large*_/λ_*small*_ = 2.0 ± 0 .1) (Figure 3E). To determine whether these actin tapering profiles were cell length-dependent, we normalized cable length by the length of the cell in which it was measured and then measured the decay lengths from these normalized profiles. Upon normalization, the actin tapering profiles collapse to a single profile with indistinguishable decay lengths (λ _*norm, small*_ = 0 .31 ± 0 .0 1, λ_*norm, large*_ = 0 .29 ± 0 .0 1) (Figure 3F), indicating that the mechanism that confers actin cable tapering is a cell length-dependent process.

### Scaling of actin cable length by tuning filament length

Our observation that cable tapering profiles depend on cell length presents two possible mechanisms by which cells can scale the length of their cables with cell length: tuning the length of filaments assembled by formins in a cell length-dependent manner (Figure 4A, Model 1), or tuning disassembly in a cell length-dependent manner (Figure 4A, Model 2). To distinguish between these two mechanisms, we conducted computational simulations and compared the simulation results to our experimental quantifications of cable length, extension rate, and tapering in smaller and larger cells.

**Figure 4:**
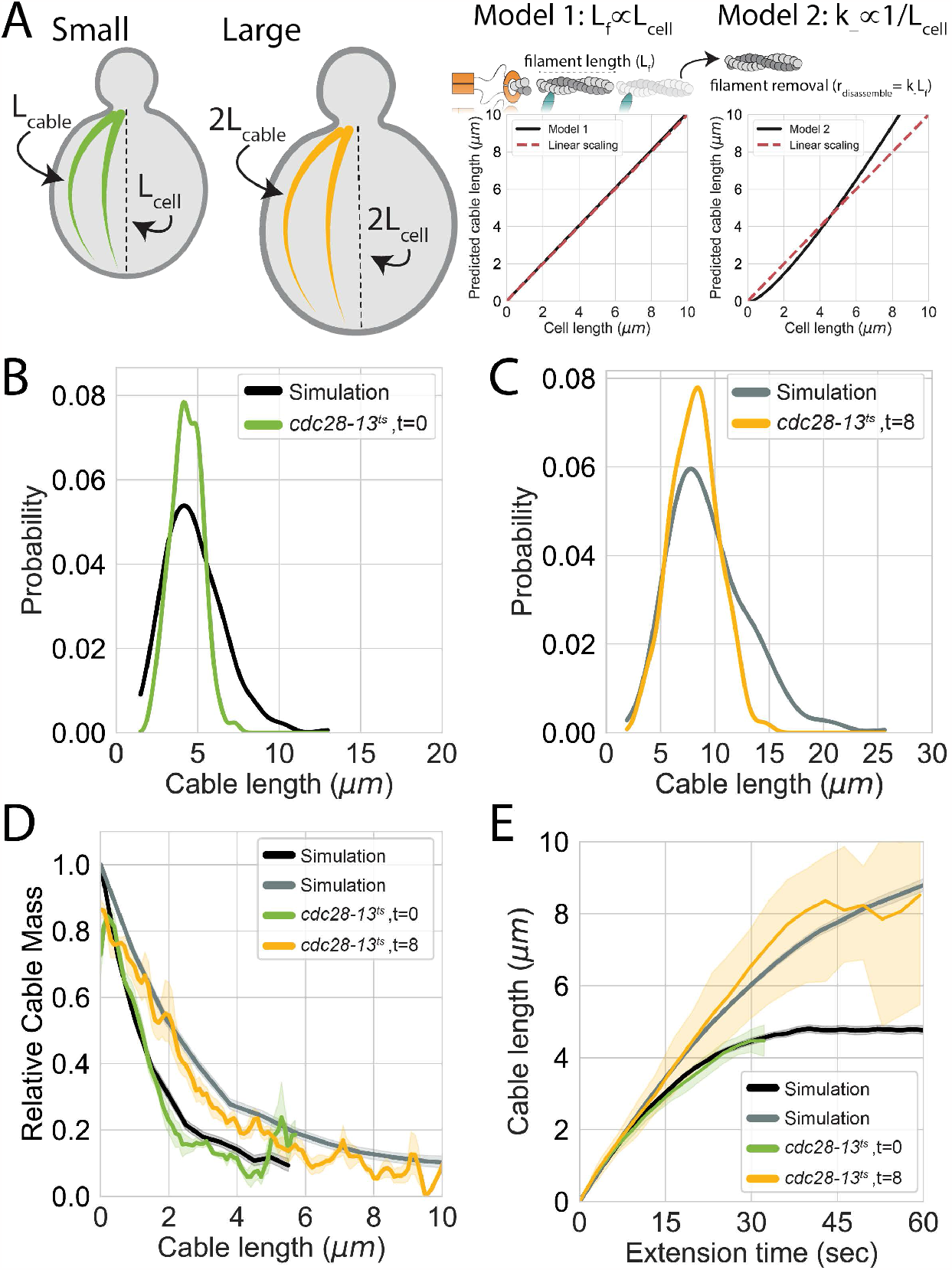
Tuning the length of formin generated filaments scales actin cable length with cell length. (A) (B-E) Comparisons between simulations conducted using the cell size specific filament lengths (black and grey lines) with experimentally measured actin cable parameters from uninduced (green lines) and induced *cdc28-13*^*ts*^ cells (yellow lines). (B-C) Comparisons of actin cable length distributions, (D) actin cable tapering profiles, and (E) actin cable extension rate. Solid lines and shading indicate mean and 95% confidence intervals, respectively.

First, we conducted simulations of cable assembly using the parameters we derived above for wildtype haploid cells, and compared these results with simulations where the disassembly rate (*k*_−_) had been scaled by cell length. We found that while the decay profiles from these simulations agree with our experimental observations (Supplemental Figure 4A), this mechanism was not able to recapitulate our other experimental observations. Specifically, the cables assembled under this mechanism were longer than expected (< *L*_*c*_ >_*large,simulation*_= 11.0 ± 1.0 μm, < *L*_*c*_ >_*large,experiment*_= 8.2 ± 0 .4μm), and that the ratio of simulated cable lengths was also greater than measured 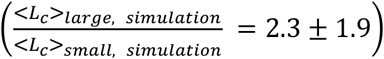 (Supplemental Figure 4B-D). Thus, it appears that tuning the disassembly rate alone cannot explain actin cable length scaling.

Next, we wanted to determine whether our experimental observations are consistent with a mechanism where the length of the filaments assembled by formins are scaled with cell length. Importantly, scaling the length of these filaments with cell length requires that both the rates of filament assembly and disassembly are also scaled in a similar manner. This is due to how these rate constants are defined in our model – each rate constant is defined by the amount of time required to either assemble or disassemble a single filament. Therefore, a 2-fold increase in filament length requires twice as much time to assemble that filament and twice as much time to disassemble that filament.

We found that our experimental data closely resemble the results of our simulations of cable assembly where the formins assemble filaments whose lengths are scaled with cell length. Specifically, there was no significant difference between mean cable length or the ratio of cable lengths between small and large cells(< *L*_*c*_ >_*large, simulation*_= 8.7 ± 0 .3μm, < *L*_*c*_ >_*small, simulation*_= 4.7 ± 0 .2μm; 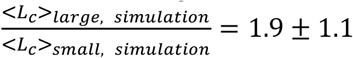; Figure 4B-C) (19). We also found that these simulations closely resemble our measurements of cable tapering (Figure 4D) and cable extension rates measured in small and large *cdc28-13*^*ts*^ cells (Figure 4E). These findings are further supported by our analytic calculations (Figure 4A for details see Supplemental Text). Thus, our experimental measurements are consistent with a mechanism where actin cable length is scaled to match cell length through a process that tunes the lengths of the filaments assembled by formins, such that formins in longer cells assemble longer filaments.

## Discussion

In this study, we present a novel, feedback-independent model of length control that describes how *S. cerevisiae* controls and scales the length of its actin cables (Figure 1C). This model differs from prior models of length control in that it does not treat each cable as a one-dimensional filament, nor does it assume that any of the model parameters are tuned in a manner that depends on cable length. Instead, our model considers the actual, two-dimensional arrangement of the cross-linked and bundled filaments that compose the cable (Figure 1C). Additionally, all processes that contribute to the assembly and maintenance of the structure (e.g., the rates of filament addition and removal, the number of nucleators) are treated as constants that are independent of the size of the structure being assembled. Despite the absence of feedback, this model recapitulates all known quantitative features of cable length control when two conditions are met: 1) the filaments that compose cables are bundled, and 2) each filament is removed from the bundle with an independent probability. Thus, rather than relying on size-dependent feedback, control over cable length instead emerges from the geometric arrangement of the shorter filaments that comprise the network.

Due to the minimal number of experimentally accessible parameters that define this model, we were able to use our quantitative experimental measurements to generate predictions for each of the parameters in our model, and then test these predictions using computational simulations. We found that our simulations of actin cable assembly using these parameters capture the key quantitative phenotypes displayed by actin cables in vivo – the distribution of cable lengths is peaked, cable extension rate decelerates as the cable grows longer, and cable thickness tapers along their length (Figure 1D-F). While the results of these simulations are very similar to our experimental measurements, we found that there are some notable differences (e.g. the width of the distribution from the simulation is greater than the width of the distribution measured experimentally). These differences between our theoretical and experimental results suggest that while our model adequately describes the mean behavior of cables (e.g., average cable length, extension rate, etc.) there are likely to be other parameters that further control the assembly and length of actin cables in vivo. Additionally, some of these differences may arise due to the complicated nature of performing quantitative experiments on such a highly dynamic cytoskeletal system. We expect that further technological developments that increase the spatial and temporal resolution with which we can observe actin cables in live cells will help to further refine the predictions for the parameters we identify in this study.

We were also interested in testing how the parameters in our model may be tuned in a cell length dependent manner to confer the scaling of actin cable length with cell length. Our prior work provided a quantitative description of how cables grow to lengths that closely match the length of the cell, however, we could only speculate about possible molecular mechanisms that would confer this behavior (19). Here, we were able use our new model of cable length control to computationally and experimentally eliminate potential mechanisms that may confer this scaling behavior.

Actin cables are assembled by two complementary sets of formins, one localized to the bud neck (Bnr1) and one localized to the bud tip (Bni1) (12–14). Our study has focused only on the cables assembled by Bnr1, which assembles and organizes cables that enter the mother cell. Prior studies have shown that Bnr1 colocalizes with components of the septin collar in regularly spaced pillars around the bud neck (30– 32). These pillars are thought to serve as sites of actin cable assembly, as actin cables have been observed to emerge from these sites as they grow into the mother cell. Additionally, it has been observed that the diameter of the bud neck scales with cell length through an unknown mechanism (33). Therefore, we sought to determine whether these sites of actin cable assembly are sensitive to changes in cell size in order to assemble longer cables in larger cells.

Our quantitative analyses of how cables are assembled in cells of different sizes revealed that while there is a greater amount of formin (Bnr1) localized to the bud neck in larger cells (Figure 2B), these cables are assembled at the same rate (Supplemental Figure 1B) and have the same initial thickness as smaller cells (Figure 3B). Additionally, we found that larger cells assemble a greater number of cables when compared with smaller cells (Figure 2E-F). Taken together, these results suggest that the molecular composition and arrangement of formins within these sites of cable assembly are likely cell size independent, but that the number of these assembly sites scales with cell length. It is currently unknown how the size, number, and composition of these cable assembly sites is determined, but we suspect that these features of the actin cable network are important to ensure that the flux of growth factors from the mother cell is sufficient to support the growth of the daughter cell. This hypothesis is supported by the observation that larger mother cells produce larger daughter cells (27, 34), and suggests that actin cables may play an important role in controlling the birth size of daughter cells.

Our analysis of actin cable tapering profiles from cells of different sizes presented two possible mechanisms to scale actin cable length with cell length – either the rate at which filaments are removed from the cable, or the length of the filaments that compose the cable are scaled with cell length. When we compared computational simulations and analytic calculations of each mechanism with our experimental measurements, we found that our data are consistent with a mechanism where the length of the filaments that compose the cable are tuned in a cell length dependent manner (Figure 4, Supplemental Text). While we have not generated direct experimental evidence to support this mechanism, prior studies have demonstrated that mutants that lack the ability to properly tune formin activity exhibit defects in actin cable length regulation and organization (16–18, 35, 36). Therefore, we suspect that the tuning of filament length may be driven by regulators that either inhibit formin activity (e.g., Smy1 and Hof1), or displace formins from the barbed ends of growing filaments (e.g., Bud14). Furthermore, it is unclear how the activity or abundance of these types of formin regulators is controlled in a cell length dependent manner. Generally, protein abundance is thought to scale with cell volume, such that their concentration is maintained across variations in cell size (37). However, recent studies have identified subsets of proteins that deviate from this behavior and either ‘sub-scale’ or ‘super-scale’ with cell volume (28, 38). Therefore, we suspect that regulators of formin activity may exhibit similar scaling behaviors, so that their abundance scales with other aspects of cell geometry (e.g., cell length or cell surface area). Alternatively, it has been recently demonstrated that cells can also exploit the different rates at which cell volume and surface area scale to tune the size of their mitotic spindle and nucleus with cell size (39, 40). Thus, it is possible that budding yeast utilize a similar mechanism to tune the activity of formins in a cell length dependent manner.

Importantly, our new model of actin cable length control was inspired by studies investigating the actin cytoskeleton arrays assembled by diverse cell types (e.g., *Listeria* and fish keratocyte lamellipodial fragments) that observed similar actin density tapering profiles (25, 26). While these structures provide fundamentally different biological functions (e.g., generating the force required for motility, or serving as tracks for intracellular transport) it appears that much of their behavior is controlled through a simple set of components – nucleators that promote the assembly of filaments, bundling or cross-linking factors that organize filaments into a higher ordered network, and disassembly factors that prune filaments from these arrays. While other studies have proposed that these diverse networks arise due to their association with specific molecular regulators, our model suggests that these higher order actin arrays have much more in common than previously thought. Furthermore, our work contributes to the emerging paradigm that, in addition to molecular regulation, the dynamics and sizes of cytoskeletal networks are encoded by their geometry (21, 41–43).

## Acknowledgments

We thank Sam Walcott, Luis Vidali, and Lishibanya Mohapatra for thoughtful comments on the manuscript.

## Methods

### Plasmids and yeast strains

All strains (see Supplemental Table 1) were constructed using standard methods. To integrate the GFP variant (Envy) at the C-terminus of the endogenous Bnr1, primers were designed with complementarity to the 3’ end of the GFP^Envy^ cassette and the C-terminal coding region of Bnr1. PCR was used to generate amplicons from the pFA6a-GFP-His3MX template that allow for selection of transformants using media lacking histidine. The parent strain, *cdc28-13*^*ts*^, was transformed with PCR products, and transformants were selected by growth on synthetic media lacking histidine. To integrate a mCherry tag at the C-terminus of the endogenous Cdc3, the plasmid pBG1533 (Cdc3-mCherry-LEU) was linearized using the restriction enzyme BglII and transformed into the parent strain, *cdc28-13*^*ts*^; *Bnr1-GFP*^*Envy*^*::His3MX*. Transformants were selected by growth on synthetic media lacking leucine.

### Induction of cell size changes

To induce increases in cell size, *cdc28-13*^*ts*^ cells were grown at the permissive temperature (25°C) overnight in synthetic complete media (SCM), then 10 µL of overnight culture was diluted into 5mL of fresh SCM. Cultures were then shifted to the restrictive temperature (37°C) for either 4 or 8 hours. After this induction, cells were returned to the permissive temperature (25°C) for one hour of growth to allow cell polarization and bud growth, and then used for imaging experiments.

### Quantitative analysis of actin cable length, number, and fluorescence intensity in fixed cells

Strains were grown at 25°C to mid-log phase (OD600 ∼ 0.3) in synthetic complete media (SCM) or were first induced for cell size changes as indicated above. Then cells were fixed in 4.4% formaldehyde for 45 minutes, washed three times in phosphate-buffered saline (1XPBS), and stained with Alexa Fluor 488-phalloidin (Life Technologies) for ≥24 hours at 4°C. Next, cells were washed three times in 1XPBS and imaged in Vectashield mounting media (Vector Laboratories). 3D stacks were collected at 0.2 μm intervals on either a Zeiss LSM 880 using Airyscan super-resolution imaging equipped with 63× 1.4 Plan-Apochromat Oil objective lens, or a Nikon Ti2-E invert confocal microscope equipped with a CSU-W1 SoRa (Yokogawa) and a Prime BSI sCMOS camera (Teledyne Photometrics) controlled by Nikon NIS-Elements Advanced Research software using a 100x, 1.45 NA objective. 3D stacks were acquired for the entire height of the cell. Airyscan image processing was performed using Zen Black software (Carl Zeiss) and SoRa image processing was performed using NIS-Elements Advanced Research software (Nikon). Quantification of actin cable length was performed as previously described (44).

To quantify actin cable number, we generated line scans of phalloidin fluorescence intensity across the approximate equator of the mother cell from background subtracted maximum intensity projection images. Lines were drawn to avoid fluorescence signal intensity associated with actin patches. Actin cables were counted by automated detection of fluorescence peaks from line scan profiles using custom Python scripts. Peaks were only identified as cables if their fluorescence intensity was greater than 20% of the maximum peak intensity within a single line scan.

To quantify the fluorescence intensity along the length of cables, we manually traced individual cables in background subtracted sum intensity projection images, from the bud neck to their terminus in the mother cell. We only included clearly discernable cables that did not intersect with other cables or actin patches. We used these line scans to record the fluorescence intensity at each position along the cable. To compare the fluorescence decay profiles of cables from different cells, the data were imported into custom Python scripts where their fluorescence intensity was normalized and rescaled so that the maximum intensity was equal to one, and the minimum fluorescence value was set to zero. These profiles were fit to a single exponential to measure their decay length.

### Simulation protocol

We used stochastic simulations to simulate the assembly of actin cables based on our two-dimensional model of cable length control. In the simulation, the system is composed of a number of rows (determined by the number of formins, N_f_, contributing to cable assembly) in which filaments of length L_f_ are added. We start these simulations with a row that contains zero filaments (i.e., a single formin that has not assembled any actin filaments) and then follow the trajectory of this row over time. For each step of the simulation, a single filament of length L_f_ is added to the row, and all other filaments within that row are selected to undergo one of the possible transitions – they are removed from the row or they remain in the row. These transitions are chosen at random based on their relative weight, which is proportional to the rate of the transition. Following these transitions, the system is updated to a new state and another step of the simulation is executed. The time elapsed between simulation steps is determined by the time required for a filament of length L_f_ to be assembled by the formin at the assembly rate, k_+_. This process is independently repeated for each row of the system, based on the number of formins (N_f_), and the length of the entire cable is determined as the distance from the initial filament position in the row to the distal end of the longest surviving filament in any row. This process is repeated for long enough time such that the length of the cable reaches steady state.

### Quantification of Bnr1 bud neck fluorescence intensity

Strains were first induced for cell size changes as indicated above and the density of each culture was measured using a spectrophotometer. The density of each culture was normalized by adding additional synthetic complete media (SCM) to the culture tube, and equal amounts of cells were harvested by centrifugation. Media was decanted and cells were resuspended in 50 µL fresh SCM and combined into a single tube and gently mixed. Approximately 5 µL of the cell suspension mixture was added onto a 1.2% agarose pad (made with SCM), and 3D stacks were collected at 0.2 μm intervals were acquired at room temperature on a Marianas spinning disk confocal system (3I, Inc, Denver,CO), consisting of a Zeiss Observer Z1 microscope equipped with a Yokagawa CSU-X1 spinning disk confocal head, a QuantEM 512SC EMCCD camera, PLAN APOCHROMAT 100X oil immersion objectives (NA 1.4) and Slidebook software. Images were processed using custom ImageJ macros. Briefly, sum intensity projections were generated and the Cdc3-mCherry channel was used for segmentation of the bud neck region of each cell. These segmentation masks were used to the measure the total fluorescence intensity of Bnr1-GFP^Envy^ for each cell and the lengths of each cell (i.e., the distance from the bud neck to the rear of the cell) were manually measured.

## Data and materials availability

Data are available in the main text or in the supplementary material. All images are archived at Zenodo and source code is available at GitHub (https://github.com/shanemc11/2DCableModel).

## Supplemental text: Two dimensional model of cable assembly

To describe the dynamics of cable assembly we consider a model which describes an actin cable as a composite structure made of actin filaments cross-linked into a bundle; see Figure 1C. Assembly of the cable proceeds at multiple formin molecules (*N*_*f*_) localized at the bud neck. We assume that each formin produces an actin filament of a fixed length (*L*_*f*_) which is incorporated into the growing actin cable at a constant rate (*k*_+_). These filaments are bundled together by crosslinkers and as a result the entire cable collectively treadmills as a single unit, extending into the mother cell at a constant extension velocity (*v*_*extension*_ = *k*_+_*L*_*f*_). In the model we describe this two-dimensional cable structure as consisting of *N*_*f*_ lanes, as shown in Figure 1C.

Once incorporated into the growing bundle, each filament has an independent probability of being targeted for removal by the action of disassembly factors. We assume that the filaments are removed at a rate *k*_−_ which makes the geometry of a cable tapered, with different lanes at any given point in time having a different length. The combined action of filament addition and removal from the *N*_*f*_ lanes leads to cable-length dynamics, where the cable length (*L*_*c*_) is defined as the length of the longest lane.

Our goal is to compute the dynamics and steady state properties of the cable length. Specifically, below we compute the probability distribution of cable lengths, its mean and variance, the steady state tapered profile of the cable, as well as the time evolution of the average cable-length, for cables that start with zero length. All these quantities we measure in our single cell experiments and, as described in the main text, we use these measurements to test our model.

### 1. Steady state cable length distribution

To compute the steady state cable length we use ideas from extreme value statistics pioneered by Fisher. To compute the probability distribution of cable lengths we consider the probability that the cable length is less than *L*:

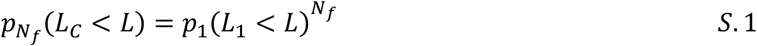

where *p*_1_(*L*_1_ < *L*) is the probability that the length *L*_1_ of one lane of the cable is less than *L*. This formula simply states that for the cable length to be less than some length *L*, then all the lanes must have a length that is smaller than *L*. The additional assumption of our model is that each lane has dynamics that are independent of every other lane, where filaments are added and removed to the lane independently of what happens to filaments in the other lanes.

To compute *p*_1_(*L*_1_ < *L*) we note that for a lane to have a length less than a specified length, the last filament in that lane must be at a distance *x* less than *L* from the formin that made it. The probability of that occurring is simply the probability that all the filaments at larger distances have been removed by the action of the disassembly factors, i.e.,

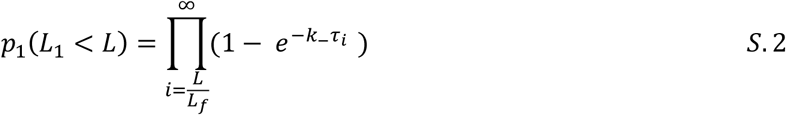

Here 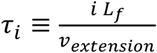 is the time that it takes the i^th^ filament to arrive at a distance *iL*_*f*_ from the formin by virtue of the whole cable structure extending at a constant speed *v*_*extension*_. The expression 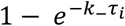is simply the probability that by time *τ*_*i*_ the i^th^ filament has been removed from the cable by disassembly factors, which remove filaments at rate *k*_−_.

Using the approximation (1 − ϵ) ≈ *e*^−ϵ^ for small ϵ, we can rewrite equation S.2 as

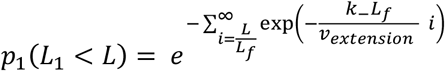

which after approximating the sum with an integral over *x* ≡ *L*_*f*_ *i* gives the formula

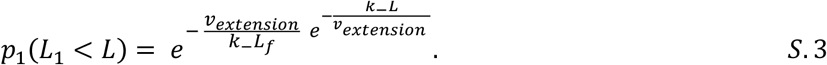

Replacing this result into equation S.1 leads to the cumulative distribution of cable lengths, when the cable consists of *N*_*f*_ lanes:

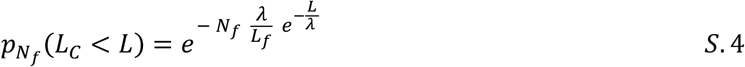

where we have introduced the characteristic length scale 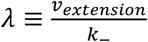 which is the average distance over which a filament is transported by the extending cable during its lifetime, which on average is 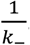. derivative of the cumulative distribution with respect to *L* yields the probability density function, which is in good agreement with stochastic simulations of the model (Supplemental Figure 2B).

Using 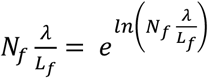 we can rewrite equation S.4 as

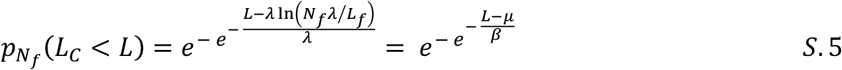

which is the Gumbel distribution with location parameter *μ* = *λ* ln(*N*_*f*_λ/*L*_*f*_) and scale parameter *β* = *λ*. The mean of the Gumbel distribution is *μ* + *βγ* (*γ* = 0.5772 … is the Euler-Mascheroni constant), while the variance is 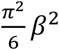. In our case, this leads to the formulas for the mean and variance of the cable length:

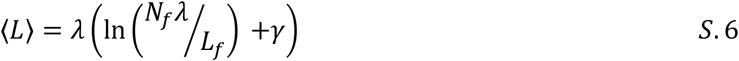

and

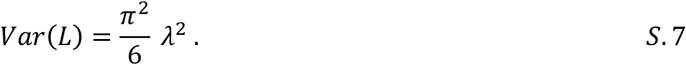

Replacing λ = *k*_+_*L*_*f*_/*k*_−_into the formulas for the mean and the variance,

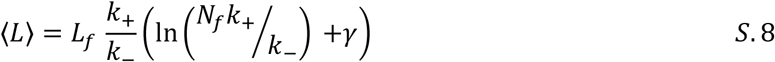

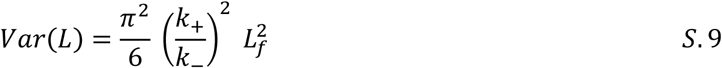

we arrive at an important result, namely that if changes in cable length are affected by changing the length of the individual filaments while keeping the number of formins and the rates of adding and removing the filaments from the cable constant, then the variance will scale as the square of the mean cable length. This is scaling we observe when changing the length of the cell, and this implies that the length of the filaments in the cable must scale with the length of the cell. This is a sharp prediction of our model that could be tested by taking EM images of cables in differently sized yeast cells.

### 2. Time evolution of the cable length

Using the model of cable assembly described above we can compute the time evolution of the average cable length, assuming that at zero time the length of the cable is zero. In experiments we obtain this quantity by watching fluorescently labeled cables extend from the bud neck to the rear of the yeast cell.

To compute the average cable length as a function of time, we start by computing the probability for the cable being shorter than some length (*L*_*c*_ < *L*) if time *t* has elapsed from the moment the cable started extending from the formins at the bud neck:

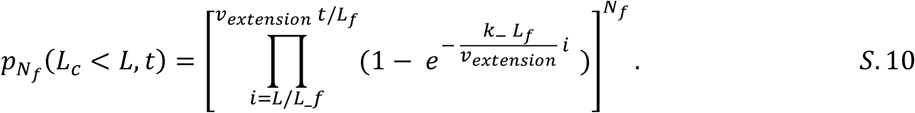

This formula assumes that the length of the cable *L* is smaller than the largest possible distance *v*_*t*_*t* that a filament can be found away from the formin, given that it has been advected with treadmilling speed *v*_*t*_ over time *t*; for larger lengths the probability is zero. The idea behind this formula is that for a cable to have a length less than *L*, then all the lanes have to be devoid of filaments that are at distances greater than *L* from the bud neck (where the formins, which inject the filaments into the cable, reside). The formula 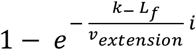 gives the probability that the filament at distance *L*_*f*_*i* (*i* is an integer that counts filaments from the bud neck) has been disassembled, given that the rate of disassembly is *k*_−_; *L*_*f*_*i* /*v*_*extension*_ is the time that filament has been in the cable since it was injected at the bud neck by the action of a formin.

Using the same approximation as in the calculation above for the steady state distribution, we can simplify equation S.10 to

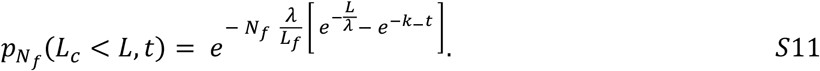

where, as above, we introduce the characteristic length scale 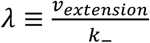, which is the average distance over which a filament is transported by the treadmilling action of the cable during its lifetime.

From the cumulative distribution, equation S.11, we can compute the mean cable length at time *t* as an integral

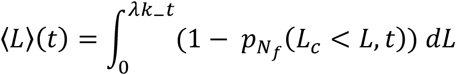

which comes out to be

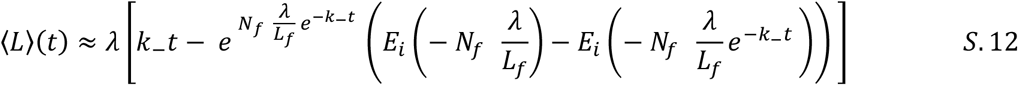

Where *E*_*i*_(*x*) is the exponential integral function. As shown in Supplemental Figure 2C in the main text this formula is in excellent agreement with stochastic simulations of the treadmilling model.

### 3. Tapering of the cable profile

Within our model we define the width of the cable *W*(*x*) as the expected number of filaments present at distance *x* away from the bud neck. Given that each cable starts out with a width that is set by the number of formins *N*_*f*_, *W*(0) = *N*_*f*_, the average number of filaments at distance *x*, is given by the survival probability that the filament was not disassembled over the time *τ* = *x*/*v*_*extension*_, where *v*_*extension*_ is the treadmilling speed of the filaments in the cable. Given that the rate of removal of filaments from the cable is *k*_−_, we find

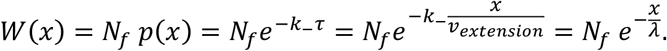

The prediction of our model is that the cable width decays exponentially with the distance away from the bud neck, which is what we observe experimentally. The decay length is set by the characteristic length λ.

## Figure legends

**Supplemental Figure 1:**
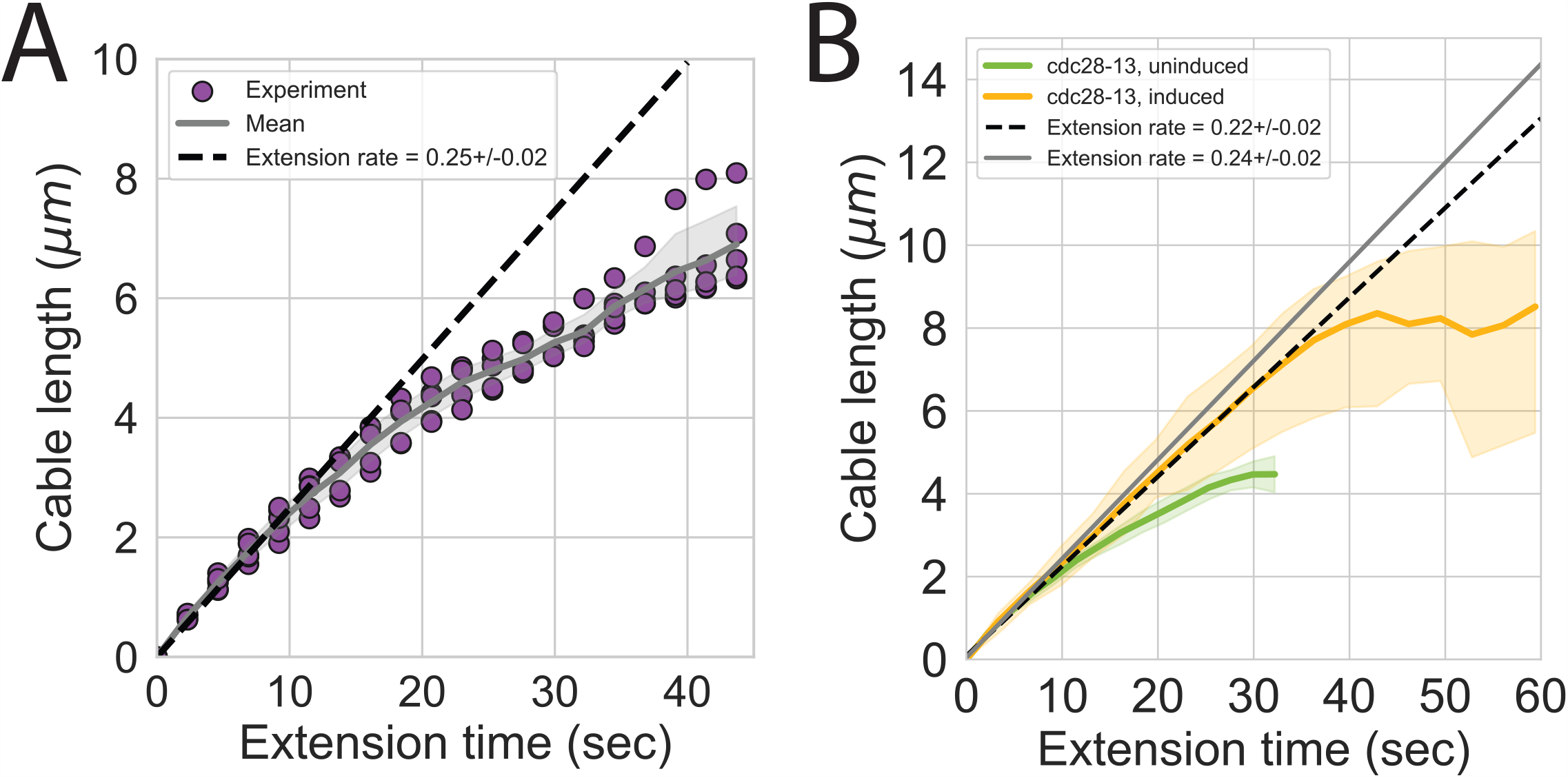
Cable extension velocity is independent of cell size. (A) Actin cable length plotted against cable extension time measured in five independent experiments (n= 82 cables). Cable extension velocity (±95%CI) (black, dashed line) was determined by linear regression using the first ∼10 seconds of extension. Symbols at each time point represent the mean for individual experiment. Solid lines and shading, mean and 95% confidence interval for all five experiments. (B) Cable extension rates for uninduced (green line) and induced *cdc28-13*^*ts*^ (yellow line) cells, from at least three independent experiments (≥57 cables/strain). Cable extension velocity (±95%CI) in uninduced (black, dashed line) and induced *cdc28-13*^*ts*^ (grey, solid line) cells was determined by linear regression using the first ∼10 seconds of extension. Solid and shading, mean and 95% confidence intervals for all experiments.

**Supplemental Figure 2:**
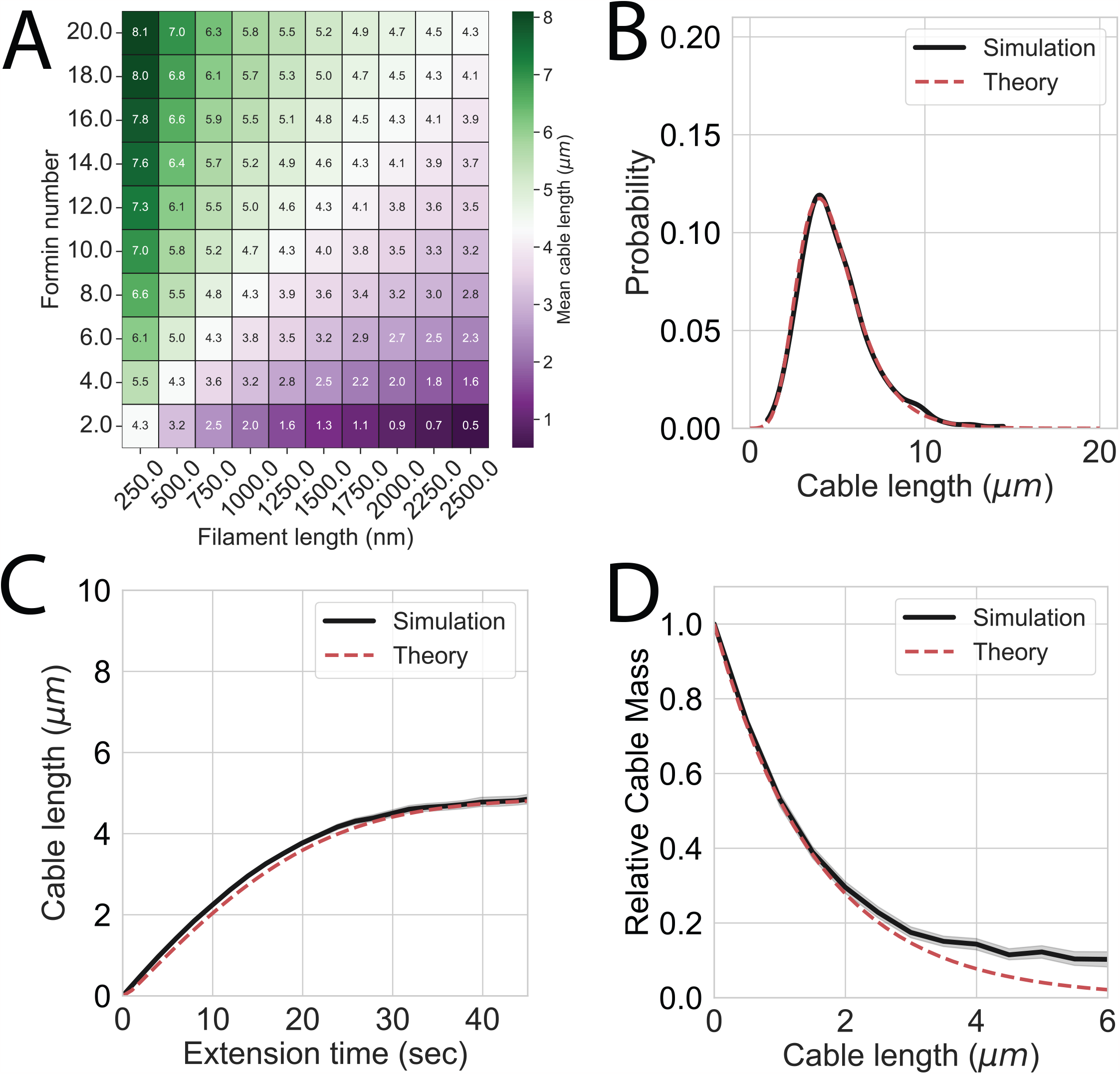
(A) Tiled heat map displaying predicted mean actin cable lengths where the number of formins and filament lengths are varied. Each tile represents the mean cable length (indicated on face of each tile) from a unique combination of formin number and filament length while λ is held constant. Divergent color coding indicates mean cable lengths that are longer (green shading) or shorter (purple shading) than mean length along the diagonal (white, < *L*_*c*_ > = 4.3 µm). (B-D) Results obtained from simulations (solid black lines) and analytical solutions (dashed red lines) show that the two-dimensional model of cable length control model produces a peaked distribution of cable lengths (A), decelerating cable extension rates (B), and cables with a tapered shape (C). The parameters used for these 1,000 independent simulations were, *k*_+_ = 0.50 sec^-1^, *k*_−_= 0.16 sec^-1^, *L*_*f*_ = 500nm, *N*_*f*_ = 4 formins. Solid lines and shading indicate mean and 95% confidence interval, respectively.

**Supplemental Figure 3:**
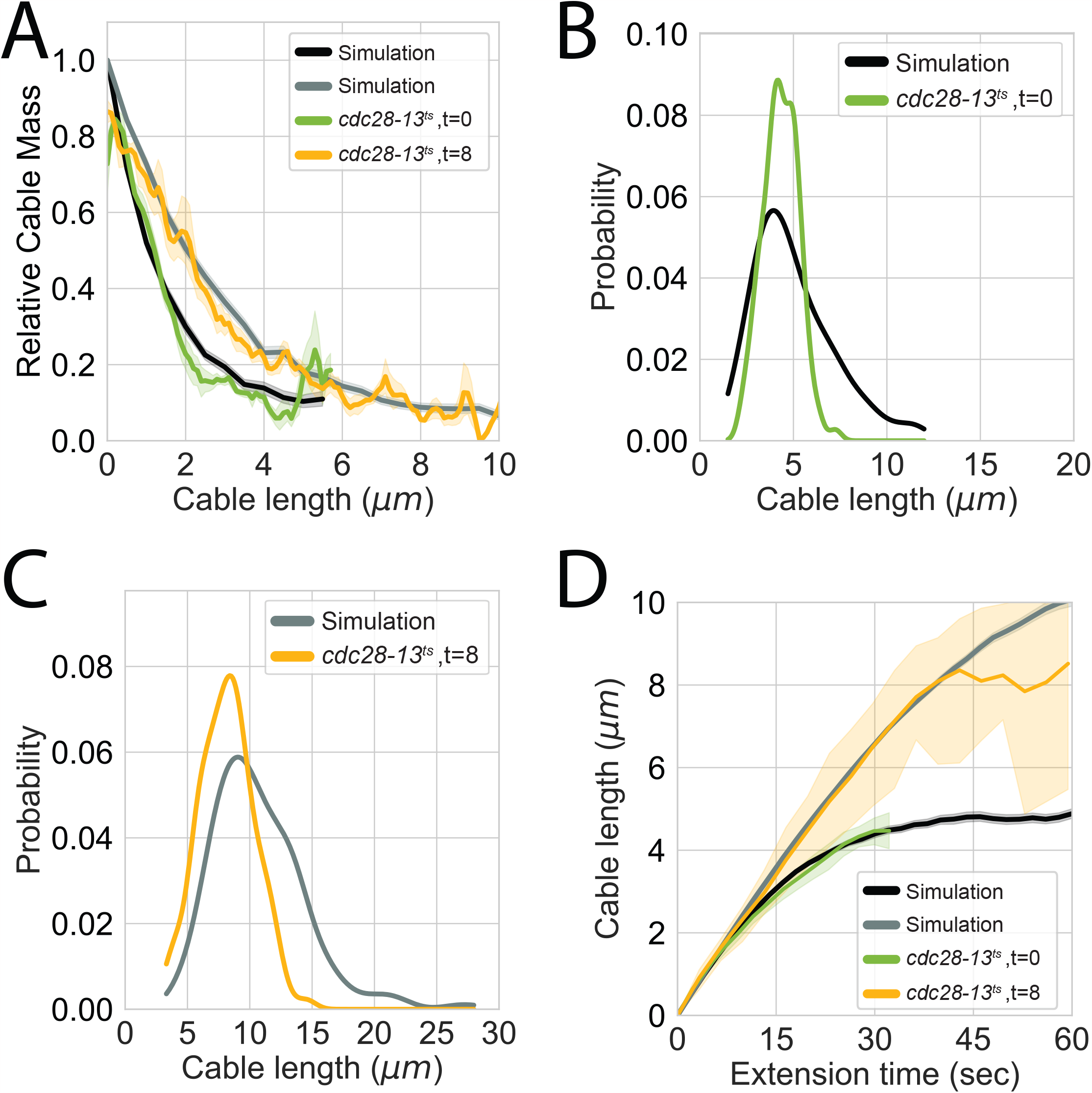
(A-D) Comparisons between simulations conducted using the cell size specific disassembly rates (black and grey lines) with experimentally measured actin cable parameters from uninduced (green lines) and induced *cdc28-13*^*ts*^ cells (yellow lines). (A) Comparisons of actin cable tapering profiles, (B-C) actin cable length distributions, and (D) actin cable extension rate. Solid lines and shading indicate mean and 95% confidence interval, respectively.

**Supplemental table 1:** Yeast strains used in this study. The genotype, source, and related data are indicated for each strain used in this study.

| Genotype | Source | Figures used in |
| --- | --- | --- |
| his3-11,15;ura3-52;leu2-3,112;ade2-1;trp1-1;psi+;ssd-;GAL+ | Lab stock | Fig. 1A-B, Fig. 1D, Fig. 1F, Fig. 2C-F |
| his3-11,15/his3-11,15;ura3-52/ura3-52;leu2-3,112/leu2-3,112;ade2-1/ade2-1;trp1-1/trp1-1;psi+;ssd-;GAL+ | Lab stock | Fig. 2E-F |
| cdc28-13; his3-11,15 trp1-1 leu2-3 ura3-1 ade2-1 | Lab stock | Fig. 2E-F, Fig. 3A-F, Fig. 4A-C, SFig. 4A-C |
| Abp140-Envy::SPHIS5; his3-11,15;ura3-52;leu2-3,112;ade2-1;trp1-1;psi+;ssd-;GAL+ | Lab stock | Fig. 1E, SFig. 3A |
| cdc28-13; Abp140-Envy::SPHIS5; his3-11,15 trp1-1 leu2-3 ura3-1 ade2-1 | Lab stock | Fig. 4D, SFig. 3B, SFig. 4D |
| cdc28-13; Bnr1-Envy::SPHIS5; Cdc3-mCherry::LEU2; his3-11,15 trp1-1 leu2-3 ura3-1 ade2-1 | This study | Fig. 3A-B |

